# Skeletal muscle MRI differentiates SBMA and ALS and correlates with disease severity

**DOI:** 10.1101/486910

**Authors:** Uros Klickovic, Luca Zampedri, Christopher DJ Sinclair, Stephen J Wastling, Karin Trimmel, Robin JMW Howard, Andrea Malaspina, Nikhil Sharma, Katie CL Sidle, Ahmed Emira, Sachit Shah, Tarek A Yousry, Michael G Hanna, Linda Greensmith, Jasper M Morrow, John S Thornton, Pietro Fratta

## Abstract

**Objective:** To investigate the use of muscle MRI for the differential diagnosis and as a disease progression biomarker for two major forms of motor neuron disorders, spinal bulbar muscular atrophy (SBMA) and amyotrophic lateral sclerosis (ALS).

**Methods:** We applied quantitative 3-point Dixon and semi-quantitative T1-weighted and STIR imaging to bulbar and lower limb muscles and performed clinical and functional assessments in ALS (n=21) and SBMA (n=21) patients, alongside healthy controls (n=16). Acquired images were analyzed for the presence of fat infiltration or edema as well as specific patterns of muscle involvement. Quantitative MRI measurements were correlated with clinical parameters of disease severity in ALS and SBMA.

**Results:** Quantitative imaging revealed significant fat infiltration in bulbar (p<0.001) and limb muscles in SBMA compared to controls (thigh: p<0.001; calf: p=0.001), identifying a characteristic pattern of muscle involvement. In ALS, semi-quantitative STIR imaging detected marked hyperintensities in lower limb muscles, distinguishing ALS from SBMA and controls. Lastly, MRI measurements correlated significantly with clinical scales of disease severity in both ALS and SBMA.

**Conclusions:** Our findings show that muscle MRI differentiates between SBMA and ALS and correlates with disease severity, supporting its use as a diagnostic tool and biomarker for disease progression. This highlights the clinical utility of muscle MRI in motor neuron disorders and contributes to establish objective outcome measures, which is crucial for the development of new drugs.

## INTRODUCTION

Amyotrophic lateral sclerosis (ALS) and spinal bulbar muscular atrophy (SBMA, also known as Kennedy’s disease) are two major motor neuron diseases. ALS is a rapidly progressive and fatal disorder characterised by relentless impairment of motor function following the degeneration of the upper and lower motor neurons (UMN, LMN)^1^. In SBMA LMN degeneration, caused by an expanded cytosine-adenine-guanine (CAG) repeat in the first exon of the androgen receptor (AR) gene^2^, induces progressive disabling bulbar and limb weakness at a slower rate^3^. At disease onset, ALS and SBMA may show similar symptoms, and distinguishing the two disorders is of paramount interest^4^.

There are still no effective disease-modifying therapies for either ALS or SBMA, and, whilst promising targets for prospective therapeutics have been identified^5^, a serious limitation for clinical trials is the shortage of sensitive and reliable outcome measures for assessing disease progression^6^.

Skeletal muscle MRI can sensitively detect muscle involvement in neuromuscular diseases^7,8^, and differentiate distinct pathological features, such as muscular fat infiltration or intra-muscular edema^9^ associated with acute denervation^10^. Muscle MRI demonstrated promise as a biomarker of disease progression in both myopathies and neuropathies^11^.

In this study we investigated muscle MRI in SBMA and ALS by studying the lower limb and the head-neck regions, which are characteristically affected in both disorders. We report that muscle MRI can differentiate between both diseases. Moreover, we show that quantitative muscle MRI measures correlate with the clinical severity of disease, thus supporting their validity as outcome measures for quantifying disease progression in clinical trials.

## METHODS

### Study Design and patient recruitment

We performed a prospective cross-sectional study, assessing muscular MRI of the head-neck region and lower limbs in 21 male adult consecutive patients with SBMA, and 21 male and female consecutive patients with ALS who were attending the national Kennedy’s disease clinic and motor neuron disease clinic at the National Hospital for Neurology and Neurosurgery, Queen Square, London, UK between 2015 and 2017. Patients with genetically confirmed mutation of androgen receptor (AR) gene were included in the SBMA group. Patients eligible for the ALS group all presented with a history of at least clinically possible disease according to revised El Escorial criteria^12^.

Additionally, 16 healthy subjects comparable to the patient groups concerning their demographic data were recruited as a control group. Differences in gender prevalence in the ALS- and SBMA-groups were taken into account, where for comparisons with ALS patients both female and male healthy volunteers were considered, whereas SBMA patients were compared to male control subjects only. General exclusion criteria for all participants were concomitant neuromuscular diseases and safety-related MRI contraindications.

Data from one ALS patient and one SBMA patient were excluded from any further analysis due to incomplete scan examination. After MRI examinations were completed, in the SBMA-group data from three patients had to be excluded due to insufficient image quality in the head-neck region. Two patients’ data were excluded for the same reason from the ALS-group. Additionally, in one control subject head-neck imaging could not be acquired due to the inability to tolerate lying still in the MR-scanner for the required examination time. The study was approved by the local ethics committee and all participants provided written informed consent at enrolment.

### Data acquisition

#### Clinical and functional testing

All participants were functionally rated using the ALS Functional Rating Scale-Revised (ALSFRS-R)^13^. In SBMA patients, functional state was additionally scored according to the SBMA Functional Rating Scale (SBMA-FRS)^14^ and supplementary measurements with the recently introduced adult myopathy assessment tool (AMAT) were performed^15^. All participants underwent detailed assessment including medical history and an examination of the clinical and neurological status.

#### MRI Imaging

MRI was performed on a clinical 3 T scanner (MAGNETOM Skyra, Siemens, Erlangen, Germany). A 20-channel head-neck matrix coil was used for imaging of the head-neck region. T1w-imaging was performed in the sagittal plane encompassing the oral cavity using a 3D isotropic fast spin-echo technique with slab selective, variable flip angle refocussing pulses (T1w-SPACE, TR = 700 ms,TE = 11.0 ms,NSA = 1, iPat = 2, 192 slices, FOV = 250 mm, voxel size: 1.0 × 1.0 × 0.9 mm^3^, total scan time of 6:04 minutes). Additionally, sagittal gradient echo 3-point Dixon sequences (TR = 125.0 ms, TE = 3.45/ 4.60/ 5.75 ms, NSA = 4, 11 slices, gap = 1 mm, FOV= 180 mm, voxel size = 0.6 × 0.6 × 10.0 mm^3^) were used to enable calculation of water- and fat-only images^16^. Total scan time for 3-point Dixon measurement was 7:54 minutes.

After this, imaging of the lower extremities was performed using the Siemens multi-channel lower-limb (‘PA matrix’) coil. Transverse 2D T1w turbo spin-echo (TSE) imaging (TR = 695 ms, TE = 9.4 ms, NSA = 1, 51 slices, gap = 3 mm, FOV = 420 mm, voxel size = 0.8 × 0.8 × 3 mm) was performed separately at the level of the mid-thigh and mid-calf, to include both limbs. Furthermore, transverse short tau inversion recovery (STIR) acquisitions (TR = 5200 ms, TE = 39 ms, TI = 220 ms, NSA = 1, iPAT = 2, 31 slices, FOV = 420 mm, voxel size = 1.1 × 1.1 × 6.0 mm, slice gap = 6 mm) were performed separately at the level of the mid-thigh and mid-calf to include both limbs. Finally, transversal T1w- gradient echo 3-point Dixon sequences comparable to the head-neck region (TR = 102.0 ms, TE = 3.45/ 4.6/ 5.75 ms, NSA = 4, 9 slices, FOV = 420 mm, voxel size = 1.3 × 1.3 × 6 mm, slice gap = 12 mm) were performed, with scan geometry adapted to the lower extremities’ regions. Total scan time for all sequences at the lower limbs was about 22 minutes.

### MRI Data Analysis

#### Semi-quantitative Muscle MRI Analysis

Semi-quantitative data analysis was based on T1w and STIR muscle imaging. Muscle fat infiltration in the thigh and calf region was assessed visually on T1w-MRI using the six-point scale (0 = normal, 1 = mild fatty streaks, 2a = early confluence, 2b = fatty infiltration 30-60%, 3 = fatty infiltration >60%, 4 = complete fat replacement) as described by Mercuri *et al.*^17^. Presence of T2w hyper-intensities at thigh and calf level were assessed visually on STIR-MRI sequences and rated by a three-point scale (0=none, 1=mild, 2=marked) as proposed by Morrow *et al.*^18^. Muscles were identified and assessed on both sides. All images were assessed and rated in consensus by two readers (U.K., K.T.) blinded to clinical diagnosis, after appropriate training.

In the head-neck region the following bulbar muscles were rated on the Mercuri scale: mastication muscles comprising m. pterygoideus medialis (Pm); m. pterygoideus lateralis (Pl); m. temporalis (TE); m. masseter (MA); swallowing muscles comprising m. buccinator (BU); m. orbicularis oris (OO); m. digastricus (Da/p - venter anterior/venter posterior); m. levator veli palatini (LvP); m. tensor veli palatini (TvP); and intrinsic and extrinsic tongue muscles comprising m. genioglossus (GG). m. geniohyoideus (GH); m. hyoglossus (HG); m. mylohyoideus (MH).

Bilaterally, lower limb muscles were assessed on both the Mercuri and Morrow scales, and included: anterior thigh compartment comprising m. rectus femoris (RF); m. vastus lateralis (VL); m. vastus intermedius (VI); m. vastus medialis (VM); posterior thigh compartment comprising m. biceps femoris (long head) and m. biceps femoris (short head) as one muscle (BFP); m. semitendinosus (ST); m. semimembranosus (SM); medial thigh compartment comprising m. adductor magnus (AM); m. sartorius (S); m. gracilis (G); anterior calf compartment comprising m. tibialis anterior (TA; including m. extensor hallucis longus); lateral calf compartment comprising. peroneus longus (PL); superficial posterior calf compartment comprising m. gastrocnemius lateralis (LG); m. gastrocnemius medialis (MG); and deep posterior calf compartment comprising the m. soleus (SO; including m. flexor digitorum longus and m. flexor hallucis longus) and m. tibialis posterior (TP).

#### Quantitative Muscle MRI Analysis

Following the semi-quantitative MRI analysis, a single observer (U.K.) blinded to subject groups defined regions of interest (ROIs) on images showing the lower limb and tongue muscles derived from the respective unprocessed shortest TE-Dixon acquisition (TE = 3.45 ms) in all subjects using the ITK-SNAP software^19^. A representative single mid-volume slice showing the previously semi-quantitatively assessed muscle groups was selected for placing the ROIs. Selection of inter-subject comparable slices was assured by identification of specific anatomical landmarks in the lower limb- (lateral tibial condyle) and the head-neck region (pituitary infundibulum). Concerning the tongue, intrinsic muscles (IM) and extrinsic muscles (GG and GH) were differentiated. After performing fat-water separation according to the method of Glover and Schneider^16^, fat fraction maps were obtained by calculating for each pixel

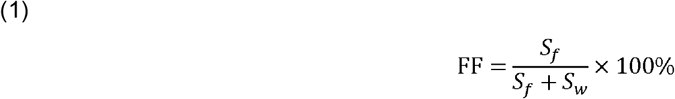

where S_f_ and S_w_ are the respective signal intensities from the same pixel location in the separated water and fat images. Muscle specific fat fraction (FF_msc_) was then obtained for each muscle and muscle compartment by overlaying the manually defined individual ROIs upon the inherently co-registered fat fraction maps^11^ and calculating the mean FF value and cross-sectional area (CSA) for each ROI.

Besides the muscle specific fat fraction (FF_msc_) for each muscle the overall muscle fat fraction percentage (FF_all_) was calculated, representing a summary measure as the mean of individual muscle values weighted by total number of voxels:

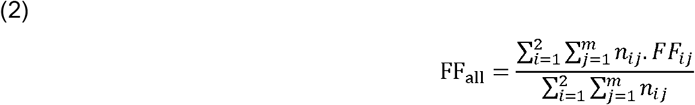

with *i*: 1=left, 2= right; *m* = total number of muscle ROIs at each level

In equation 2 *n*_*ij*_ denotes the number of voxels in a certain muscle specific ROI of the left or right side, where index j relates to a respective muscle on each side. Analogously, the MRI-based functional remaining muscle area (fRMA_msc_), defined as the CSA of the muscle tissue not replaced by fat, was computed as:

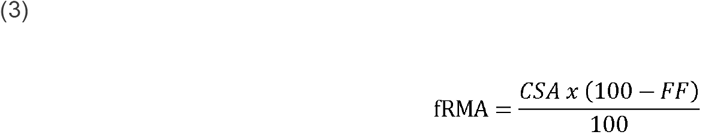

#### Statistical Analysis

Statistical analyses were performed with SPSS version 22 (Armonk, NY) with an alpha level of 0.05. As appropriate to the distribution of data, measures are reported as mean ± SD respectively median and interquartile range (IQR). For inter-group comparisons, Kruskal-Wallis-tests, 2-sample t-tests and Mann-Whitney-U-tests were applied as appropriate. Missing data were excluded from analyses. Correlations of MRI data with clinical measures were investigated with Spearman (rho) or Pearson coefficients as appropriate for the distribution of data.

## RESULTS

### Participant demographics and clinical findings

The study included two patient groups, SBMA (N = 21) and ALS (N = 21), along with healthy controls (N = 16). For analysis exclusion criteria we refer to the methods section. Mean age was 50.7 (SD 17) years and 54.4 (SD 14.6; p = 0.53) years in the SBMA and SBMA-control group respectively (**Table 1**). Mean age was 57.3 (SD 14.8) y and 55.4 (SD 13.5; p = 0.69) years in the ALS and ALS-control group respectively. Height and weight did not significantly differ between the two patient groups and controls, except for ALS patients having a significantly lower body mass index compared to their control group (p=0.009). In both patient groups, ALSFRS-R and its lower limb (LL) subscale scores were similar (ALSFRS-R ‒ total score, SBMA: 42 (31-48), ALS: 41 (28–47); ALSFRS-R ‒ LL subscore, SBMA: 5 (2–8), ALS: 5.5 (3–8); ALSFRS-R ‒ bulbar subscore, SBMA: 10.5 (9–12), ALS: 12 (5–12) and significantly reduced compared to healthy controls (SBMA vs. CTR; p < 0.001, ALS vs. CTR; p < 0.001; **Table 1**).

**Table 1.**
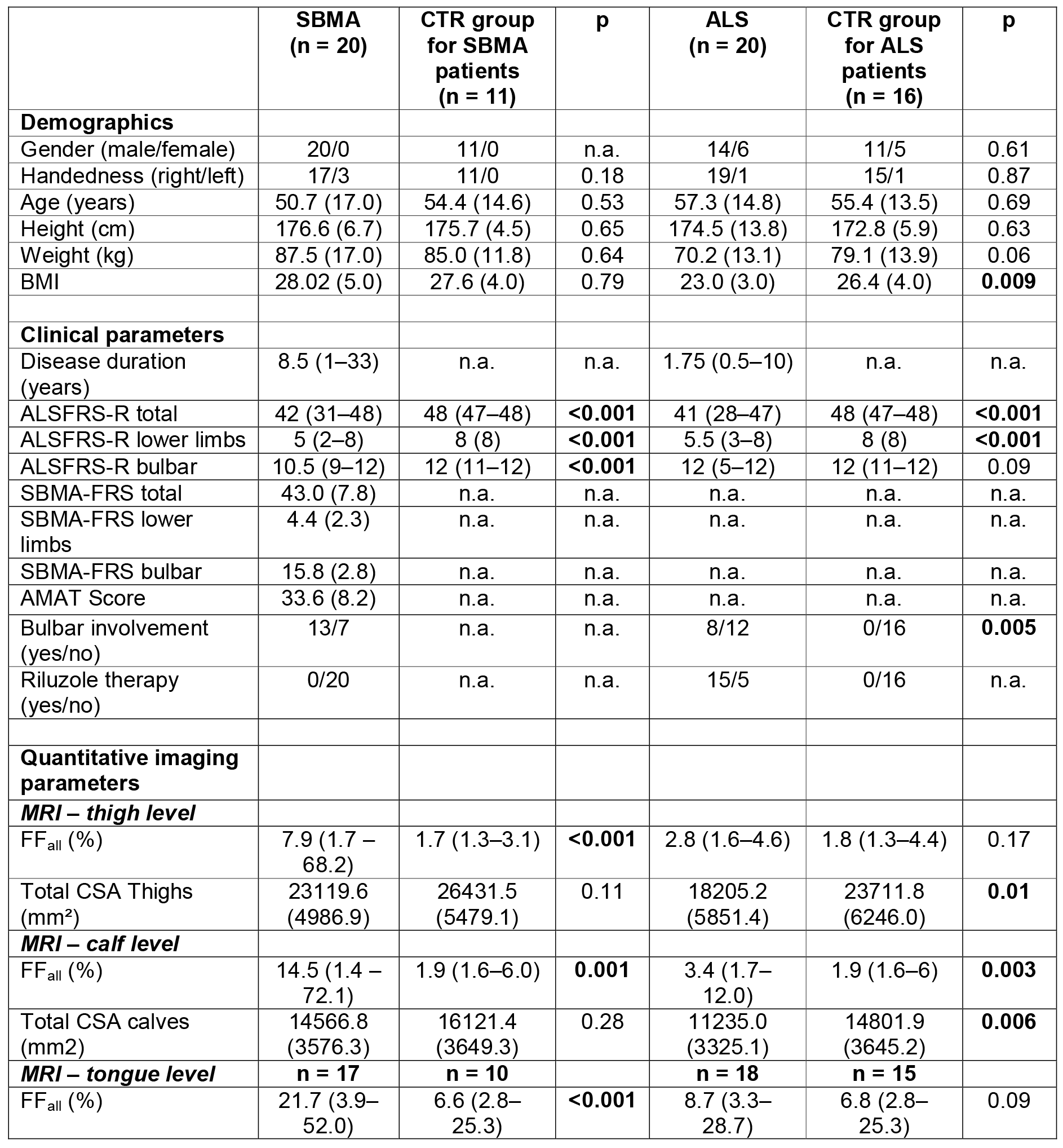
Demographic and clinical data of SBMA patients, ALS patients and controls. Data are presented as mean and SD or median and range as appropriate to distribution of the data. ALS: amyotrophic lateral sclerosis, ALSFRS-R: ALS functional rating scale ‒ revised, AMAT: adult myopathy assessment tool, BMI: body mass index, CSA: cross sectional area, CTR: controls, FFall: overall muscle fat fraction percentage, MRI: magnetic resonance imaging, SBMA: spinal and bulbar muscular atrophy, SBMAFRS: SBMA functional rating scale.

### Thigh muscle fat infiltration differentiates SBMA from ALS and defines specific patterns of muscle involvement

To assess the pattern and the severity of muscle fat infiltration in SBMA and ALS, we acquired T1w^10^ and quantitative 3-point Dixon^11^ MRI from LL muscles. We rated the T1w images according to the Mercuri scale^17^. Whilst whole-thigh SBMA Mercuri scores were significantly increased compared to controls (median 2, IQR 1 vs median 1, IQR 0; M-W-U = 10, p < 0.001), there was no difference in ALS compared to controls (median 1, IQR 0 vs median 1, IQR 0; M-W-U = 130, p = 0.35; **Figure 1A**). We identified a previously unrecognised pattern of muscle involvement in the SBMA patient group, affecting both anterior and posterior thigh muscle compartments (TMC) with relative sparing of the medial TMC. Specifically, the vastus lateralis (VL) and the semimembranosus (SM) were the most severely affected muscles in the anterior and posterior TMC, whilst in the medial TMC no muscle showed severe involvement (Mercuri grade 4). In contrast to SBMA, no ALS patient showed severe muscle fat involvement by scoring Mercuri grade 3 or 4 in any of the TMC (**Figure 1A**).

**Figure 1.**
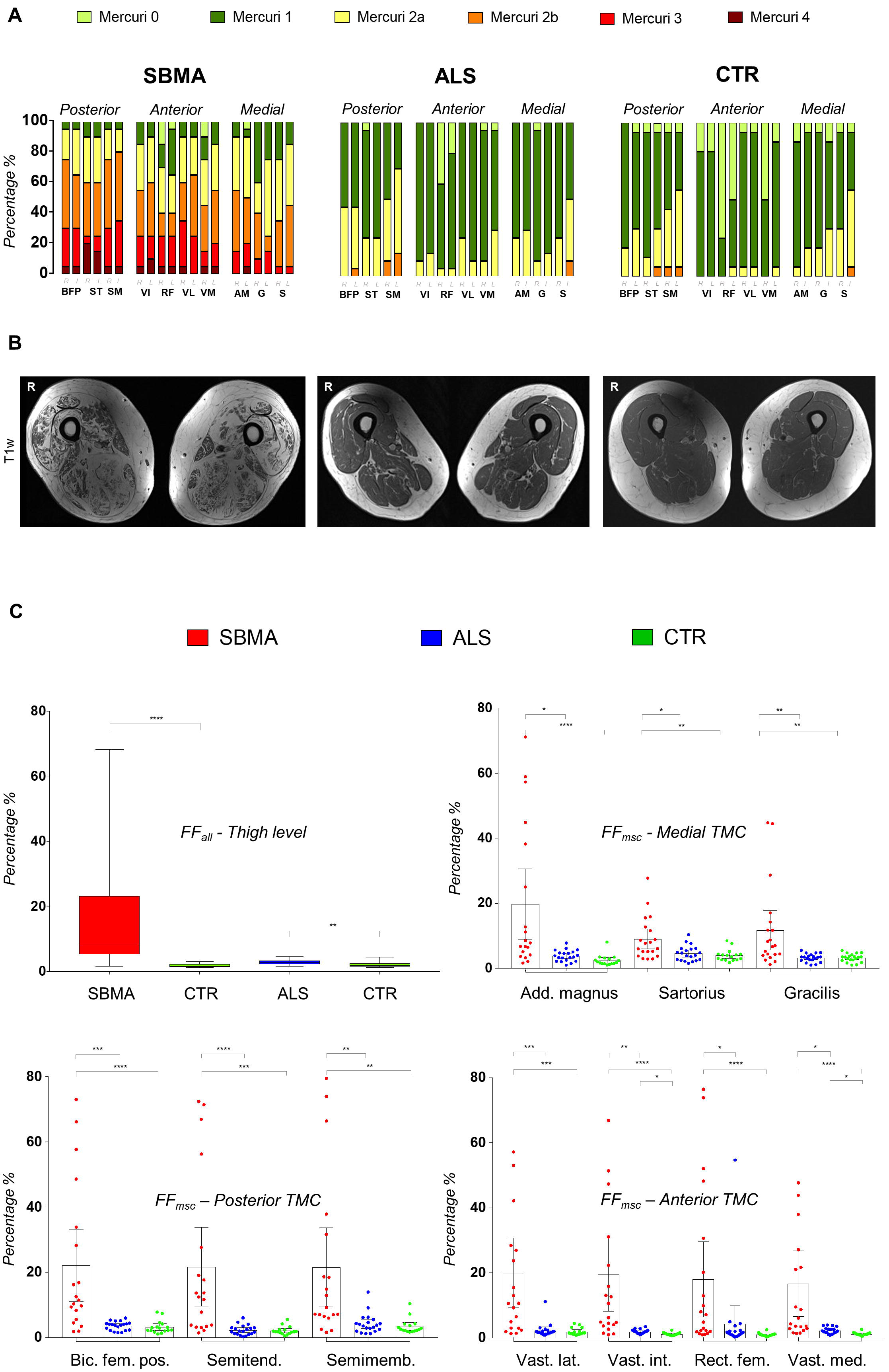
Semiquantitative and quantitative muscle MRI analysis: thigh muscle compartment. **(A)** The proportion of Mercuri scores of thigh muscle compartments (TMC) in all study groups *(for muscle abbreviations see methods section).* SBMA, Spinal and bulbar muscular atrophy; ALS, amyotrophic lateral sclerosis; CTR, control subjects. Mercuri scores are higher in SBMA group compared to CTR group (M-W-U = 10, p < 0.001). **(B)** Corresponding axial T 1w sample images of the right and left thigh in SBMA (left), ALS (middle) and CTR (right). **(C)** Overall muscle fat fraction percentage (FF_all_) at thigh level in SBMA, ALS and CTR (upper row left). Boxes represent median and CI, whiskers show range. FF_all_ is significantly increased at thigh level in SBMA and ALS group compared to their age-matched controls (SBMA vs. CTR: M-W-U = 12, p < 0.001; ALS vs. CTR: M-W-U = 75, p = 0.006). Muscle specific fat fraction (FF_msc_) of medial (upper row right), posterior (lower row left) and anterior (lower row right) right TMC. Data are shown as dot plots with boxes representing mean and whiskers representing 95% CI. Asterisks indicate p values of post-hoc pairwise comparisons between study groups of Kruskal-Wallis test results for each right thigh muscle. * = 0.05; ** = 0.005; *** 0.0005; **** < 0.0001. Add. magnus, adductor magus; Bic fem pos, biceps femoris posterior; Semitend, semitendinosus; Semimeb, semimembranosus; Vat lat, vastus lateralis; Vast int, vastus intermedius; Rect fem, rectus femoris; Vast med, vastus medialis.

Overall muscle fat fraction percentage (FF_all_) at mid-thigh level was significantly higher in both patient groups compared to their matched control group (SBMA vs CTR: median 7.9%, IQR 17.93% vs. median 1.67%, IQR 0.85%; M-W-U = 12; p < 0.001; ALS vs CTR: median 2.79%, IQR 1.13% vs. median 1.79%, IQR 0.98%; M-W-U = 75; p = 0.006; Figure 1C). The muscle specific fat fraction (FF_msc_) confirmed the pattern identified in the semi-quantitative assessments: in SBMA patients, the highest FF_msc_ was observed in the posterior TMC (biceps femoris posterior (BFP), followed by semitendinosus (ST) and SM) and in the anterior TMC (VL, followed by vastus intermedius (VI) and rectus femoris (RF)); the medial TMC was relatively spared with the lowest FF_msc_ in SBMA patients (**Figure 1C**). Despite a significantly higher overall FF_all_ compared to their matched controls, no specific pattern of fat infiltration was observed on single muscle analysis in ALS patients. However, there was significant atrophy of thigh muscles in ALS patients compared to controls (**Table 1**).

In summary, widespread intramuscular fat accumulation was observed in SBMA patients occurring with a preferential involvement of muscles in the anterior and posterior TMC.

### Calf muscle fat infiltration differentiates SBMA from ALS and defines specific patterns of muscle involvement

Mercuri scores for calf T1w images were significantly altered in both SBMA and ALS patients compared to their matched controls (SBMA vs. CTR: median 3, IQR 1.88 vs. median 1, IQR 0; M-W-U = 10, p = 0.001; ALS vs. CTR: median 1,5, IQR 1 vs. median 1, IQR 0; M-W-U = 130, p = 0.008). SBMA patients showed a previously non-described pattern of muscle involvement with relative sparing specifically of the tibialis anterior (TA) and posterior (TP) muscles and predominant involvement of the superficial and deep posterior calf muscle compartments (CMC), with more than 50% of patients scoring Mercuri grade 3 or 4 in gastrocnemius medialis (MG), soleus (So) and gastrocnemius lateralis (LG) muscles (**Figure 2A**). Conversely, none of the ALS patients or healthy controls had Mercuri grade 3 or 4 in any of the CMC. However, more than 70% of ALS patients showed moderate fatty degeneration (Mercuri grade 2a or 2b) in the lateral CMC, affecting mostly the peroneus longus (PL).

**Figure 2.**
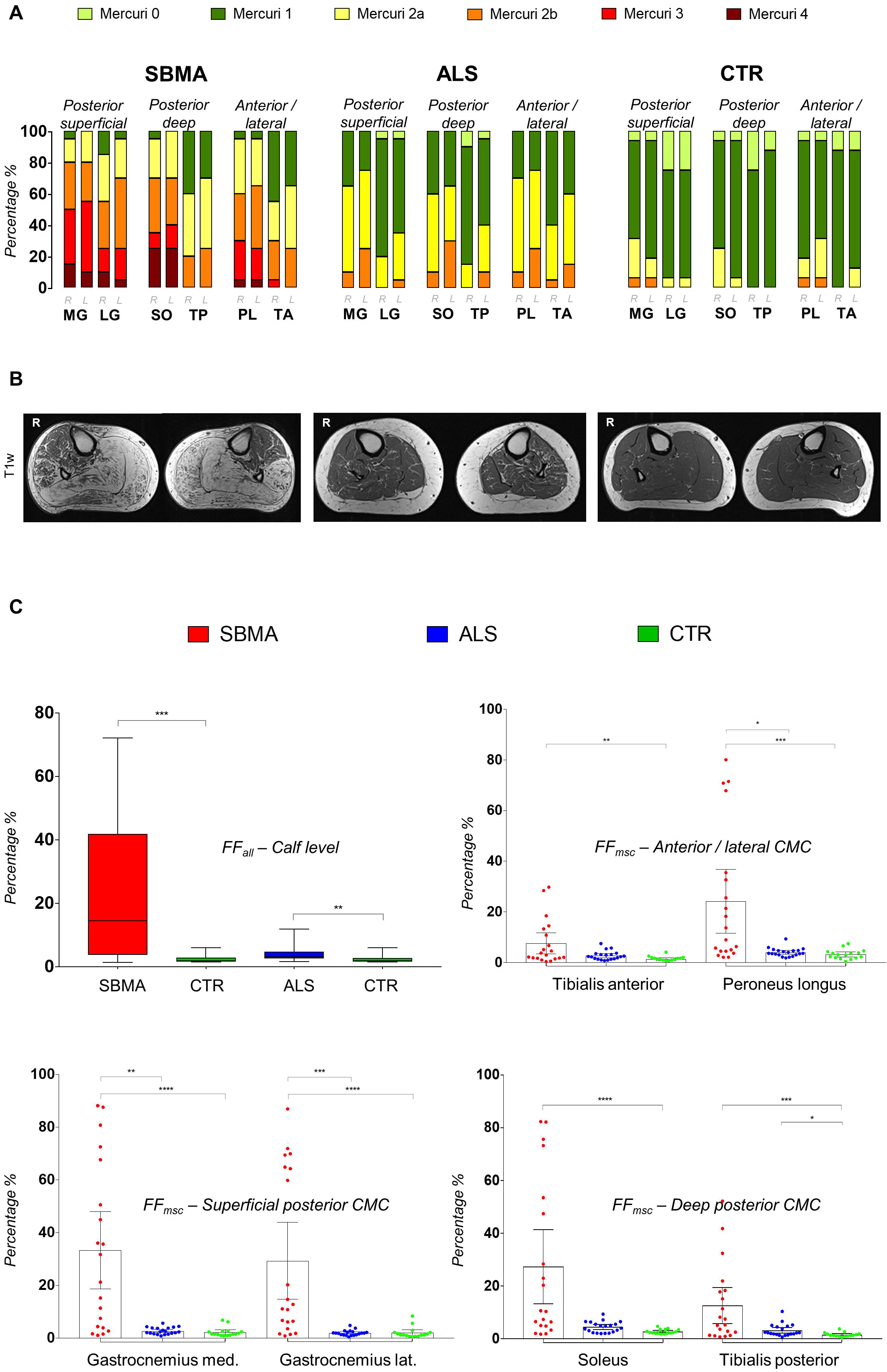
Semiquantitative and quantitative muscle MRI analysis: calf muscle compartment. **(A)** The proportion of Mercuri scores of calf muscle compartments (CMC) in all study groups *(for muscle abbreviations see methods section).* SBMA, Spinal and bulbar muscular atrophy; ALS, amyotrophic lateral sclerosis; CTR, control subjects. Mercuri scores are higher in both patient groups compared to their matched CTR group (SBMA vs. CTR: M-W-U = 10, p = 0.001; ALS vs. CTR: M-W-U = 130, p = 0.008). **(B)** Corresponding axial T1w sample images of the right and left calf in SBMA (left), ALS (middle) and CTR (right). **(C)** Overall muscle fat fraction percentage (FFall) in SBMA, ALS and CTR (upper row left). Boxes represent median and CI, whiskers show range. FF_all_ is significantly increased at calf level in SBMA and ALS group compared to their age-matched controls (SBMA vs. CTR: M-W-U = 31, p = 0.001; ALS vs. CTR: M-W-U = 69, p = 0.003). Muscle specific fat fraction (FF_msc_) of anterior/lateral (upper row right), superficial posterior (lower row left) and deep posterior (lower row right) right CMC. Data are shown as dot plots with boxes representing mean and whiskers representing 95% CI. Asterisks indicate p values of post-hoc pairwise comparisons between study groups of Kruskal-Wallis test results for each right calf muscle. * = 0.05; ** = 0.005; *** 0.0005; **** < 0.0001. MED, medial; LAT, lateral.

Quantitatively, FF_all_ were higher in both patient groups compared to controls, albeit with much lower levels than in ALS (SBMA vs CTR: median 14.54%, IQR 38.17% vs median 1.87%, IQR 1.23%; M-W-U = 31; p = 0.001; ALS vs CTR: median 3.34%, IQR 2.07% vs. median 1.92%, IQR 1.08%; M-W-U = 69; p = 0.003; **Figure 2C**). Analysis of the FF_msc_ confirmed the semi-quantitative findings of predominant posterior CMC affection and sparing of TA and TP. At calf level, the highest FF_msc_ were found in the superficial posterior CMC. Although FF_msc_ of specific calf muscles were generally lower in ALS than SBMA patients, in keeping with semi-quantitative findings, the lateral CMC showed the highest FF_msc_ within ALS patients (**Figure 2C**). Compared to controls, significant atrophy of calf muscles in ALS patients was observed (**Table 1**).

In summary, widespread intramuscular fat accumulation, occurring with a specific pattern relatively sparing the TA and TP muscles and more severely affecting the posterior compartment muscles, was observed in the SBMA patients. ALS also showed changes, albeit at lesser levels, and with a different pattern, the lateral compartment being the most affected.

### Fat suppressed STIR images show significant changes in both SBMA and ALS

As the rapid course of ALS may contribute to the modest level of fat infiltration in this condition, we investigated whether semi-quantitative fat suppressed STIR sequences would reveal more identifiable changes deriving from quickly developing muscle denervation. Marked STIR hyperintensities were observed in almost all TMC and CMC in both patient groups compared to their age-matched controls.

Using the rating scale proposed by Morrow *et al.*^18^, marked muscle tissue hyperintensities (Morrow grade 2) were detected in the anterior and posterior TMC of SBMA patients (**Figure 3A**); in the calves of SBMA patients, STIR abnormalities were observed in all muscle compartments (**Figure 3C**). In the ALS patient group, the anterior TMC was the most altered, albeit signal abnormalities were detected in every CMC. In healthy controls, no marked hyperintensities were observed in any of the TCM, whilst in the calf, we noticed STIR signal hyperintensities of MG, previously described as “central stripe”^10^, which corresponds to the muscle end-plate region of this muscle (**Figure 3C, 3D**).

**Figure 3.**
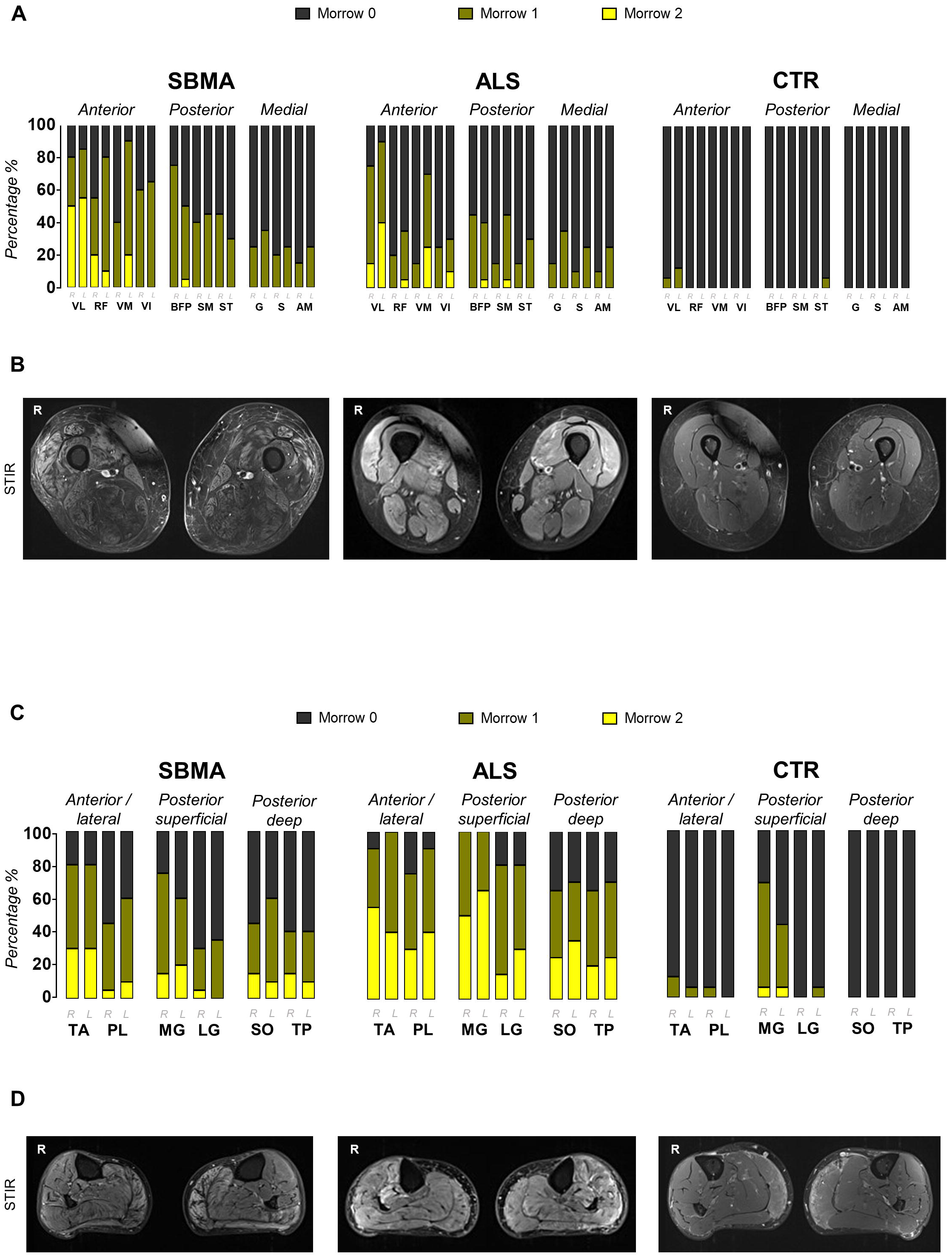
Semiquantitative muscle MRI analysis: STIR imaging. **(A)** The proportion of Morrow scores of thigh muscle compartments (TMC) in all study groups *(for muscle abbreviations see methods section)*. SBMA, Spinal and bulbar muscular atrophy; ALS, amyotrophic lateral sclerosis; CTR, control subjects. STIR hyperintensities were observed in both patient groups. No marked signal abnormalities were observed in healthy controls. **(B)** Corresponding sample STIR axial images of the right and left thigh in SBMA (left), ALS (middle) and CTR (right). **(C)** The proportion of Morrow scores of calf muscle compartments (CMC) in all study groups *(for muscle abbreviations see methods section)*. STIR hyperintensities were observed in both patient groups. Apart from MG, no STIR hyperintensities were observed in controls. **(D)** The corresponding sample STIR axial images of the right and left calf in SBMA (left), ALS (middle) and CTR (right).

In summary, differently from T1w sequences, STIR sequences detect changes in both conditions, with alterations in the calves being more marked in ALS.

### Muscle MRI detects widespread muscle changes at bulbar level in SBMA

Although bulbar muscles are affected in both SBMA and ALS, these have rarely been included in previous muscle MRI studies^20,21^. We therefore assessed fatty changes of bulbar muscles using both T1w and 3-point Dixon sequences.

T1w imaging showed a moderate to severe involvement (Mercuri grade 2b – 4) of mastication and swallowing muscles in SBMA, whilst in contrast, ALS patients showed predominantly only mild to moderate muscle fat infiltration of these muscle groups (Mercuri grade 1 – 2b; **Figure 4A**). Both patient groups showed moderate to severe involvement (Mercuri 2b or higher) of intrinsic and extrinsic tongue muscles, including the genioglossus (GG), geniohyoideus (GH), hyoglossus (HG) and mylohyoideus (MH). Notably, 12.5% of healthy subjects also showed moderate muscle fat infiltration (Mercuri grade 2b) in extrinsic tongue muscles (**Figure 4A**). Quantitative FF_all_ were significantly higher in SBMA patients (median 21.69%, IQR 16.66%), compared to their matched controls (median 6.63, IQR 3.50; M-W-U = 18; p < 0.001; Figure 4B). FF_msc_ were highest in the intrinsic tongue muscles, followed by extrinsic tongue muscles GG (median 16.56%, IQR 16%) and GH (median 10.29%, IQR 15%) in SBMA patients. No significant difference in FF_all_ was seen between ALS patients (median 8.68%, IQR 5.74%) and their matched controls (median 6.76%, IQR 2.84%; M-W-U = 94; p = 0.15; **Figure 4B**).

**Figure 4.**
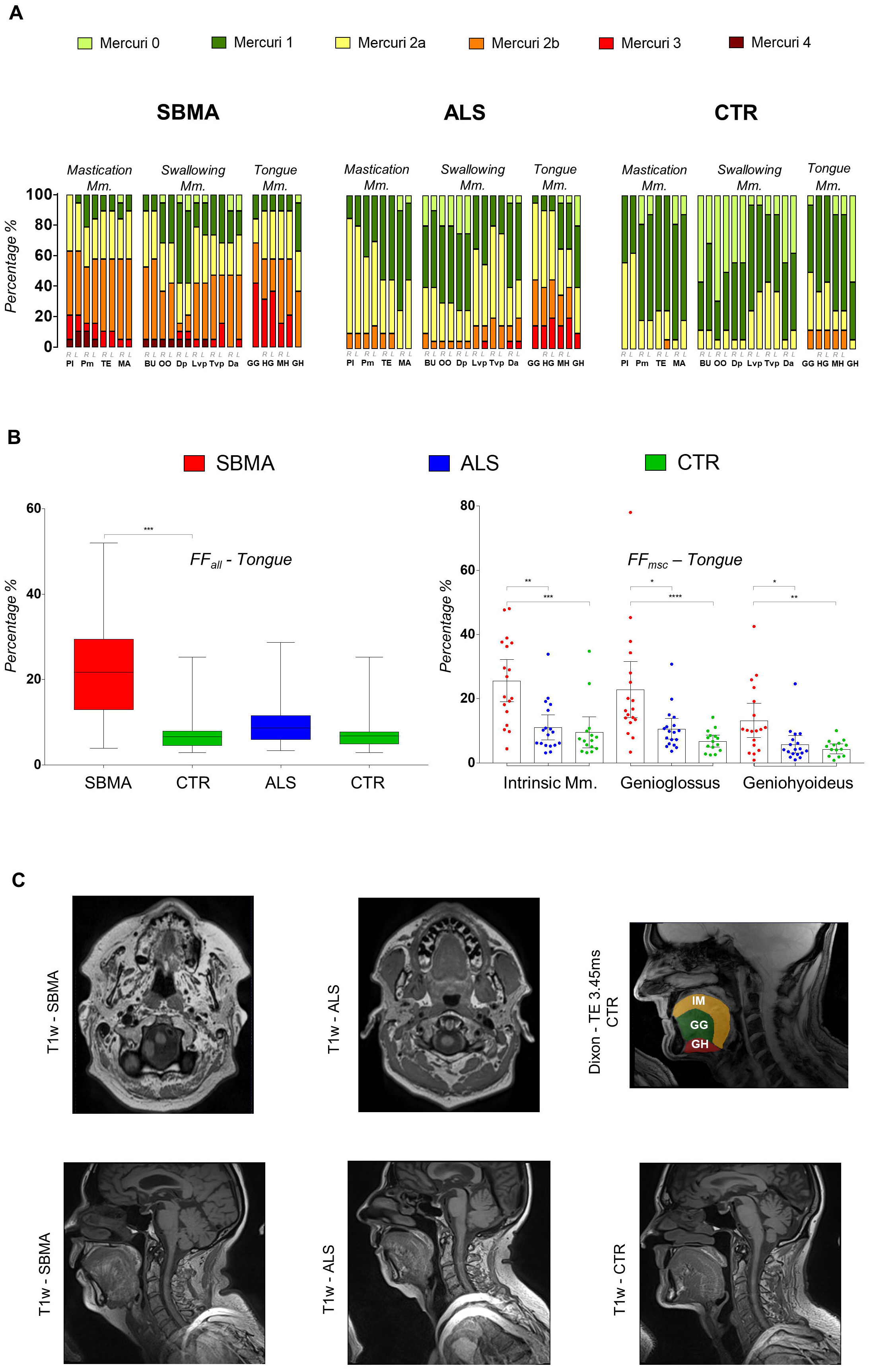
Semiquantitative and quantitative muscle MRI analysis: head and neck region. **(A)** The proportion of Mercuri scores of bulbar muscles in all study groups *(for muscle abbreviations see methods section).* SBMA, Spinal and bulbar muscular atrophy; ALS, amyotrophic lateral sclerosis; CTR, control subjects. Mercuri scores of bulbar muscles are higher in SBMA group and ALS group compared to their matched CTR group (SBMA vs CTR: M-W-U = 21, p < 0.001; ALS vs. CTR: M-W-U = 90, p = 0.03). **(B)** Overall muscle fat fraction percentage (FF_all_) of tongue muscles in SBMA, ALS and CTR (left). Boxes represent median and CI, whiskers show range. FF_all_ is significantly higher in SBMA compared to controls (M-W-U =18, p < 0.001). Muscle specific fat fraction (FF_msc_) of intrinsic and extrinsic (genioglossus and geniohyoideus) tongue muscles (right). Asterisks indicate p values of post-hoc pairwise comparisons between study groups of Kruskal-Wallis test results for each tongue muscle. * = 0.05; ** = 0.005; *** 0.0005; **** < 0.0001. **(C)** Corresponding T1w sample images of head and neck muscles of SBMA (upper row left) and ALS (upper row middle). Regions of interest (ROI) of intrinsic and extrinsic tongue muscles derived from the unprocessed shortest TE - Dixon sequence (TE = 3.45 ms) of CTR (upper row right). Sagittal T1w sample images of tongue muscles of SBMA (lower row left), ALS (lower row middle) and CTR (lower row right). Mm, muscles; IM, intrinsic muscles; GG, Genioglossus; GH, Geniohyoideus.

In summary, fat infiltration in bulbar muscles is able to differentiate SBMA from both ALS and controls.

### Muscle MRI correlates with functional rating scales in MND

In order to test whether the MRI findings reflected disease severity, we calculated correlations between muscle MRI measures and functional measures, using established clinical rating instruments for each condition: the ALSFRS-R for ALS patients and the SBMA-FRS and the AMAT score for SBMA.

We initially asked whether the overall mean FF_all_ of combined thigh and calf level correlated with the LL subscale from ALSFRS-R and found there was a strong negative correlation between FFall and disability in SBMA patients (ρ = −0.86, p < 0.001) and a significant negative correlation also in ALS patients (ρ = −0.47, p = 0.04; **Figure 5A**). In SBMA, the FF_all_ strongly negatively correlated with the total AMAT score (ρ = −0.77, p < 0.001. **Figure 5C**). It is important to note that FF_all_ did not correlate with age in healthy controls (ρ = −0.06, p = 0.8).

**Figure 5.**
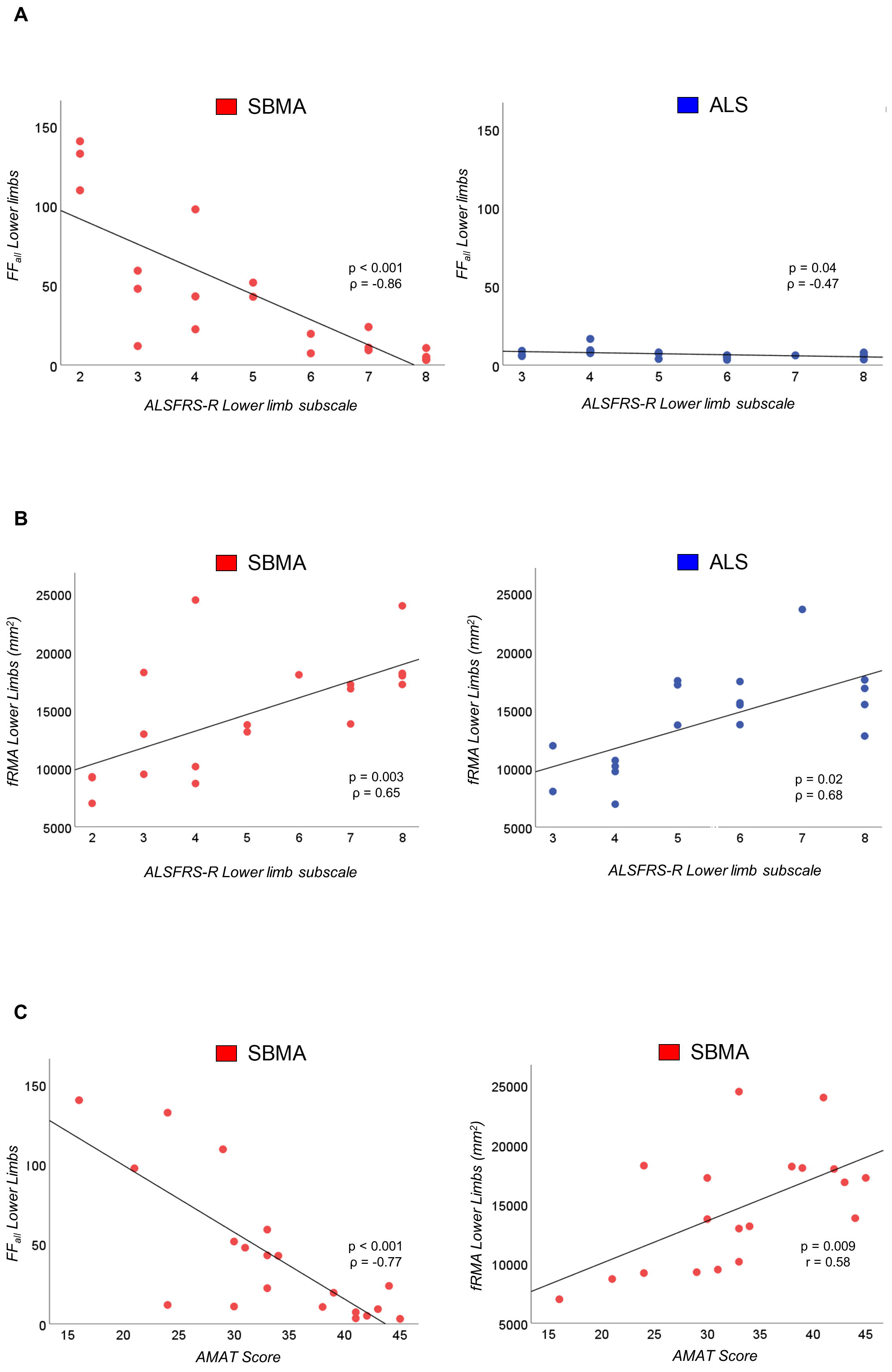
Correlation of muscle MRI parameters with functional rating scales. **(A)** FF_all_ of LL shows a significant correlation with ALSFRS-R LL subscale in both patient groups (left; SBMA: ρ = −0.86, p < 0.001; right: ALS: ρ=−0.47, p = 0.04). **(B)** fRMA of LL shows a significant correlation with ALSFRS-R LL subscale in both patient groups (left; SBMA: p = 0.65, p = 0.003; right: ALS: ρ = 0.68, p = 0.02). **(C)** FF_all_ of LL correlates significantly with AMAT score in SBMA patients (left: ρ = −0.77, p < 0.01). Significant correlations were observed between fRMA of LL and AMAT score in SBMA patients (right, r = 0.58, p = 0.009). LL, lower limbs; AMAT, adult myopathy assessment tool; ALSFRS-R, ALS functional rating scale ‒ revised; FF_all_, overall muscle fat fraction percentage; fRMA, functional remaining muscle area.

Since our results show that fat infiltration is not a prominent feature in ALS, whilst active denervation changes are occurring, we next asked whether assessment of the functional remaining muscle area (fRMA_msc_) would take muscle atrophy into account and provide a better measure for ALS. Indeed, in ALS patients the fRMA_msc_ correlated strongly with ALSFRS-R LL subscales (ρ = 0.68, p = 0.002; **Figure 5B**). The combined thigh and calf level fRMA_msc_ of SBMA patients also correlated strongly with SBMA-FRS LL subscales (ρ = 0.70, p = 0.003), and with the AMAT score (r = 0.58, p = 0.009, **Figure 5C**).

In summary, MRI measures correlate with functional measures in both SBMA and ALS, supporting their exploration as quantitative biomarkers of disease progression.

## DISCUSSION

In the present study, we performed skeletal muscle MRI imaging on cohorts of two major forms of adult motor neuron disease, ALS and SBMA. Furthermore, for the first time, we also extended quantitative fat fraction quantitation protocols to the head and neck region, given the involvement of bulbar muscles in these diseases. The results of our study have important clinical relevance; they identify a novel pattern of muscle involvement and show that quantitative and semi-quantitative skeletal muscle MRI can differentiate ALS from SBMA. In addition, they support the validity of muscle MRI as a disease-progression biomarker, as we show that MRI-quantified muscle FF and fRMA correlate with clinical measures of both diseases.

Muscle MRI studies in ALS^22,23^ and SBMA patients^24^ have mostly been limited to a small number of cases and have not performed quantitative assessments^25^. Recently, Dahlqvist et al. published results further supporting the validity of Dixon imaging in SBMA^26^. We have studied cohorts of significant size, using qualitative and quantitative MRI methods previously validated in other neuromuscular disease patient groups. Importantly, whilst a focussed analysis of the lower limbs was appropriate for such conditions, here we also developed reproducible protocols demonstrating the feasibility and potential value of extending MRI investigations to bulbar muscles, which are crucially involved in ALS and SBMA. While electromyography (EMG) is a rather restricted technique due to its invasive nature and requires complete relaxation of the tongue for appropriate recording^27^, muscle MRI can be used to investigate specific anatomical regions at the bulbar level.

Our results suggest that different muscle changes occur in ALS and SBMA, with SBMA showing marked fat infiltration on T1w MRI, whilst in ALS, increased edema seen on STIR images is the most prominent feature. Muscle denervation is a common pathological feature in ALS^28^, and the increase in extracellular fluid within the denervated muscle^29^ is sensitively reflected by hyperintensities on T2-weighted fat-suppressed MRI sequences as STIR. Using whole-body MRI, Jenkins *et al.* recently reported higher relative T2 muscle signals in ALS compared to controls, which correlated with clinical weakness and lower motor unit number^30^. In accord with these findings, we observed marked STIR hyperintensities in all thigh and calf muscles in both SBMA and ALS patients, highlighting the potential of fat-suppressed MRI sequences to reveal changes deriving from muscle denervation in the rapid course of disease progression in ALS.

Furthermore, we have identified consistent patterns of muscle involvement and sparing in SBMA. Thus, fat accumulation was observed predominantly in the posterior muscle compartment at thigh and calf level (with relative sparing of the medial thigh compartment and anterior calf compartment), and both intrinsic and extrinsic muscles at tongue level. SBMA and ALS are in differential diagnosis and approximately 13% of SBMA patients have been reported to have previously been given a diagnosis of ALS^31,32^. Our findings support the use of muscle MRI in the diagnostic work-up of patients with prevalent lower motor neuron involvement and suspected MND.

Importantly, we here also show that the MRI-obtained fRMA_msc_ correlates with established clinical parameters of disease progression in both diseases. Functional RMA takes into account both the FF_msc_ of muscles, which is crucial in SBMA, and the muscle atrophy which is typical of ALS. The ability of MRI changes to reflect clinical involvement support their future use as disease progression biomarkers. Indeed, there is currently an intensive search for biomarkers of disease progression in MND in order to carry out more effective clinical trials^6^. SBMA is slowly progressive and rare, and more reliable, quantitative outcome measures would both reduce the duration as well as the size, and cost, of clinical trials. Survival has often been used in ALS trials^33^, but novel sensitive outcome measures in ALS would also enable shorter trials to be undertaken, reducing costs and allowing more therapies to be tested. Muscle MRI appears to fulfil the criteria for such a biomarker, due to its reproducibility, its capability of generating quantitative measurements and its observer independence^34^. Furthermore, fat infiltration or denervation-related changes detected by muscle MRI may provide the ground for the development of neurochemical markers, which reflect these changes. However, further studies will be necessary to assess the sensitivity of muscle MRI to detect progressive changes in these diseases and to determine the full potential of muscle MRI as a reliable outcome measure for ALS and SBMA.

## ACKNOWLEDGEMENTS

We thank all study participants and families for their participation to research that has made this work possible. This study was supported by the UCLH NIHR Biomedical Research Centre, Kennedy’s Disease UK (KD-UK) and the Institute of Neurology Kennedy’s Disease Research Fund. UK is funded by KD-UK. PF is funded by an MRC CS LEW Fellowship and by the UCLH NIHR Biomedical Research Centre. LG is the Graham Watts Senior Research Fellow funded by the Brain Research Trust.

---

| Name | Location | Role | Contribution |
| --- | --- | --- | --- |
| Uros Klickovic MD, MRes | UCL Queen Square Institute of Neurology, University College London, London, UK | Author | Data acquisition and data analysis; statistical analysis; writing of the manuscript. |
| Luca Zampedri | UCL Queen Square Institute of Neurology, University College London, London, UK | Author | Data acquisition and data analysis. |
| Christopher DJ Sinclair PhD | Neuroradiological Academic Unit, UCL Queen Square Institute of Neurology, University College London, London, UK | Author | Data acquisition and data analysis. |
| Stephen J Wastling PhD | Neuroradiological Academic Unit, UCL Queen Square Institute of Neurology, University College London, London, UK | Author | Data acquisition and data analysis. |
| Karin Trimmel MD, PhD | UCL Queen Square Institute of Neurology, University College London, London, UK | Author | Data acquisition and data analysis. |
| Robin JMW Howard MD | UCL Queen Square Institute of Neurology, University College London, London, UK | Author | Data acquisition and data analysis. |
| Andrea Malaspina MD, PhD | Blizard Institute, Queen Mary, University of London, London, UK | Author | Data acquisition and data analysis. |
| Nikhil Sharma MD, PhD | UCL Queen Square Institute of Neurology, University College London, London, UK | Author | Data acquisition and data analysis. |
| Katie CL Sidle MD | UCL Queen Square Institute of Neurology, University College London, London, UK | Author | Data acquisition and data analysis. |
| Ahmed Emira | Neuroradiological Academic Unit, UCL Queen Square Institute of Neurology, University College London, London, UK | Author | Data acquisition and data analysis. |
| Sachit Shah MD | Neuroradiological Academic Unit, UCL Queen Square Institute of Neurology, University College London, London, UK | Author | Data acquisition and data analysis. |
| Prof Tarek A Yousry MD | Neuroradiological Academic Unit, UCL Queen Square Institute of Neurology, University College London, London, UK | Author | Critical revision of the manuscript. |
| Michael G Hanna FRCP | UCL Queen Square Institute of Neurology, University College London, London, UK | Author | Critical revision of the manuscript. |
| Linda Greensmith PhD | UCL Queen Square Institute of Neurology, University College London, London, UK | Author | Critical revision of the manuscript. |
| Jasper M Morrow MD, PhD | UCL Queen Square Institute of Neurology, University College London, London, UK | Author | Design and conceptualized study; data acquisition and data analysis; critical revision of the manuscript. |
| John S Thornton PhD | Neuroradiological Academic Unit, UCL Queen Square Institute of Neurology, University College London, London, UK | Author | Design and conceptualized study; data acquisition and data analysis; critical revision of the manuscript. |
| Pietro Fratta MD, PhD | UCL Queen Square Institute of Neurology, University College London, London, UK | Author | Design and conceptualized study; data acquisition and data analysis; statistical analysis; writing of the manuscript. |

**APPENDIX 1 - AUTHOR CONTRIBUTIONS**

